# Fusion detection and quantification by pseudoalignment

**DOI:** 10.1101/166322

**Authors:** Páll Melsted, Shannon Hateley, Isaac Charles Joseph, Harold Pimentel, Nicolas Bray, Lior Pachter

**Affiliations:** Faculty of Industrial Engineering, Mechanical Engineering, and Computer Science, University of Iceland; Department of Molecular and Cell Biology, University of California, Berkeley; Graduate Program in Computational Biology, University of California, Berkeley; Department of Computer Science, University of California, Berkeley; Innovative Genomics Initiative, University of California, Berkeley; Division of Biology and Biological Engineering, California Institute of Technology, Pasadena

## Abstract

RNA sequencing in cancer cells is a powerful technique to detect chromosomal rearrangements, allowing for *de novo* discovery of actively expressed fusion genes. Here we focus on the problem of detecting gene fusions from raw sequencing data, assembling the reads to define fusion transcripts and their associated breakpoints, and quantifying their abundances. Building on the pseudoalignment idea that simplifies and accelerates transcript quantification, we introduce a novel approach to fusion detection based on inspecting paired reads that cannot be pseudoaligned due to conflicting matches. The method and software, called pizzly, filters false positives, assembles new transcripts from the fusion reads, and reports candidate fusions. With pizzly, fusion detection from raw RNA-Seq reads can be performed in a matter of minutes, making the program suitable for the analysis of large cancer gene expression databases and for clinical use. pizzly is available at https://github.com/pmelsted/pizzly

---

Accelerated acquisition of heterogeneous, somatic mutations across cells within a tumor or pre-tumorous tissue is a widespread occurrence in most cancer types^1^. Resultant mutations are sufficient to induce tumorigenesis and drive tumor progression and metastasis through activation of proto-oncogenes and deactivation of tumor suppressor genes. The accumulation of mutations can be induced through the malfunction of several biological pathways, involving several corresponding mutation classes^2^. One common and particularly deleterious class of mutation is chromosomal rearrangement. Chromosomal rearrangements typically occur when two or more double-strand breaks are incorrectly repaired through rearrangement, deletion, or duplication. Large-scale changes to gene expression from such rearrangements are associated with a number of disorders^3^. When the rearrangement breakpoints are at gene regulatory and/or coding regions, the disruption can result in a deregulated or a *chimeric* fusion gene (Figure 1). Gene fusions are associated with many types of cancer and play a major role in tumorigenesis4,5,6,7,8,9.

**Figure 1:**
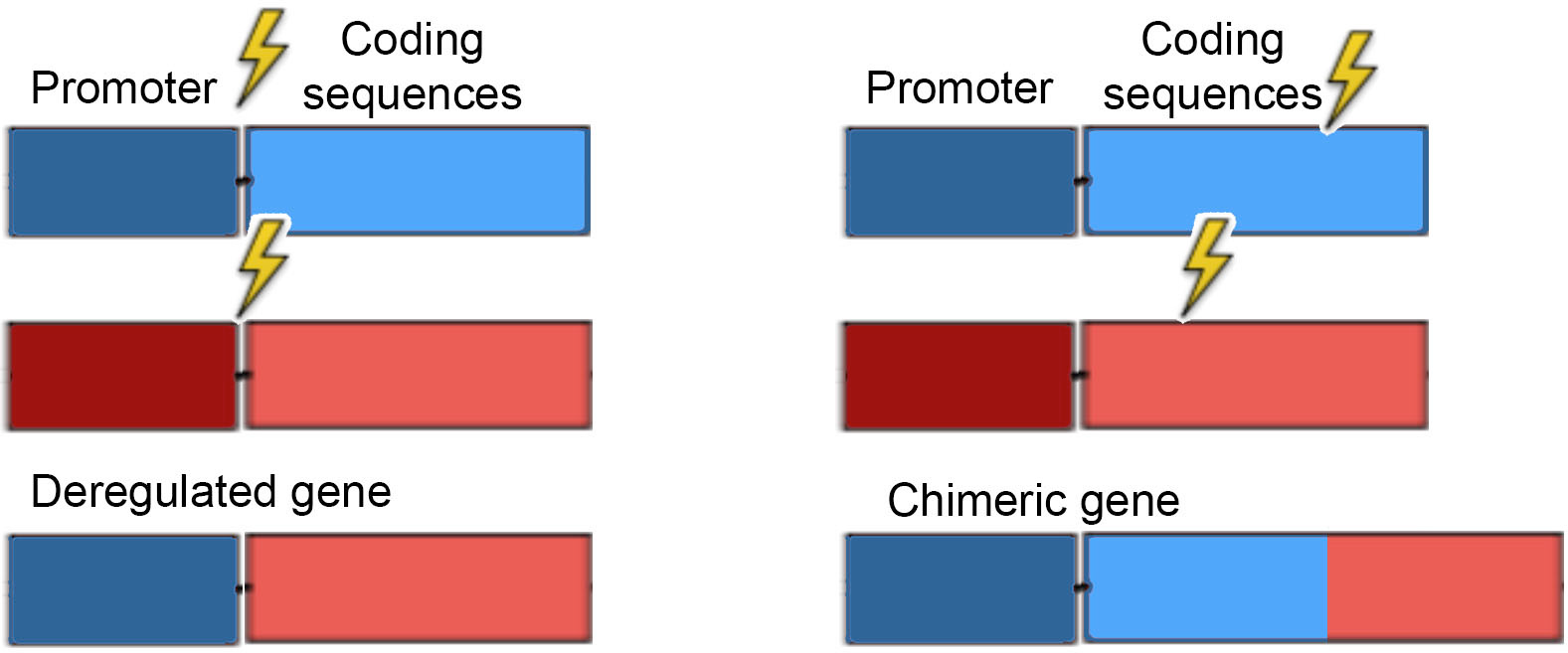
Fusion genes resulting from chromosomal rearrangements. If the chromosomal rearrangement is made at coding and/or regulatory regions, a deregulated or chimeric gene can occur. Lightening bolt indicates chromosomal breakpoint.

Currently, most fusion detection methods rely upon RNA sequencing^10^ (RNA-Seq) because it is cost-effective, covers all transcribed regions of the genome, and is highly interpretable. Several computational tools to detect fusions from RNA-Seq reads have been developed, and these typically operate via three steps: First, they detect “spliced” reads where the first part of a (possibly paired-end) read maps to one gene and the second part maps to another gene. Second, reads are aggregated into candidate fusion junctions through realignment-based grouping. Finally, candidate fusions are filtered based on a number of heuristics^11^.

In practice, identification of gene fusions via RNA-Seq is difficult and error-prone. While over 20,000 gene fusions have been identified in the three major neoplasia subtypes, fewer than a 1,000 of them have been confirmed as recurrent^3^. Factors contributing to the inflated number of positive fusions include incorrect read mapping, transplicing, and template switching. Incorrect mapping is often a result of fusion detection tools attempting to map fusions from transcripts arising from repetitive regions of the genome such as germline segmental duplications. Transplicing can induce read-through events, where transcription continues across genomically adjacent genes to be transcribed into one RNA transcript. Template switching events also contribute to the count of false positive gene fusions as the RNA polymerase jumps from one DNA template to another during transcription^12^. These produce low but detectable baseline levels of fusion genes in wild-type as well as cancerous cells, but are usually not of interest in cancer sequencing efforts as they are not causally involved in tumorigenesis, cancer progression, or metastasis^13^.

To filter out false positives, fusion detection tools employ a set of heuristics based upon the contributing biological factors just described, as well as a number of ad hoc filters. The result is that existing fusion detection tools each contain some mixture of the following disadvantages: they may be computationally demanding, display poor sensitivity/specificity, be difficult to install, questionable heuristics may bias predictions, they may not resolve breakpoints, and may fail to quantify fusions.

Here, we introduce an improved method for gene fusion discovery utilizing the publicly-available software kallisto^15^ which is based on pseudoalignment, along with a downstream processing tool, pizzly, that (1) filters false positives using biologically-relevant heuristics, (2) assembles transcripts at breakpoint resolution, and (3) quantifies fusion abundances. This method is computationally efficient allowing for laptop-based analysis, and enables consistent investigation of large cancer gene expression databases to probe for tumor-specific and cancer-type agnostic recurrent fusions.

## Methods

Fusion junctions are detected using a two-stage method. The first stage is implemented in kallisto and detects individual reads or read pairs, whose constituent parts pseudoalign to a reference transcriptome but that in combination fail to pseudoalign. pizzly takes the output of the potential fusion junctions found by kallisto and performs a detailed analysis of the associated reads by aligning them across the putative junctions. Additionally, pizzly is annotation aware, i.e. it uses information about the genomic coordinates and gene identities of each isoform to identify possible false positives arising from repetitive sequences across the genome.

### kallisto Fusion Stage

The kallisto program was enhanced to include an option to search for reads that could support fusion junctions. kallisto uses a *k*-mer based index for the reference transcriptome when computing a pseudoalignment for reads. For each *k*-mer, the index records the set of transcripts containing this *k*-mer, which is its equivalence class (EC). For a normal read arising from a single transcript, all matching k-mers that are supported by at least that transcript and the intersection of the ECs for the read will be non-empty and contain the true transcript. For reads or read pairs that span a fusion junction, the ECs from each side of the junction will have an empty intersection and thus be discarded from further consideration by kallisto. When run in fusion finding mode, kallisto identifies read pairs whose intersection of ECs is empty, yet for which some of the k-mers matched. Such reads are reported as fusion output when either one of the following holds: (1) each read has a non-empty EC intersection separately, but combined the intsersection is empty (this is consistent with Figure 2 “paired fusion reads,” where the reads come from opposite sides of the fusion junction) or (2) one of the reads can be split into two parts such that the first part of the read has a non-empty EC intersection and the remainder of the read, along with the other read from the pair, has a non-empty EC intersection (this is consistent with Figure 2 “split fusion reads,” where one of the reads spans the fusion junction). When a potential split of the reads has been identified, kallisto checks all matching k-mers and requires that the union of ECs on either side is empty. This is to lower the number of false positives due to reads from unannotated transcripts that resemble fusions between related transcripts. All read pairs matching these criteria are saved along with supporting information about the matching transcripts.

**Figure 2:**
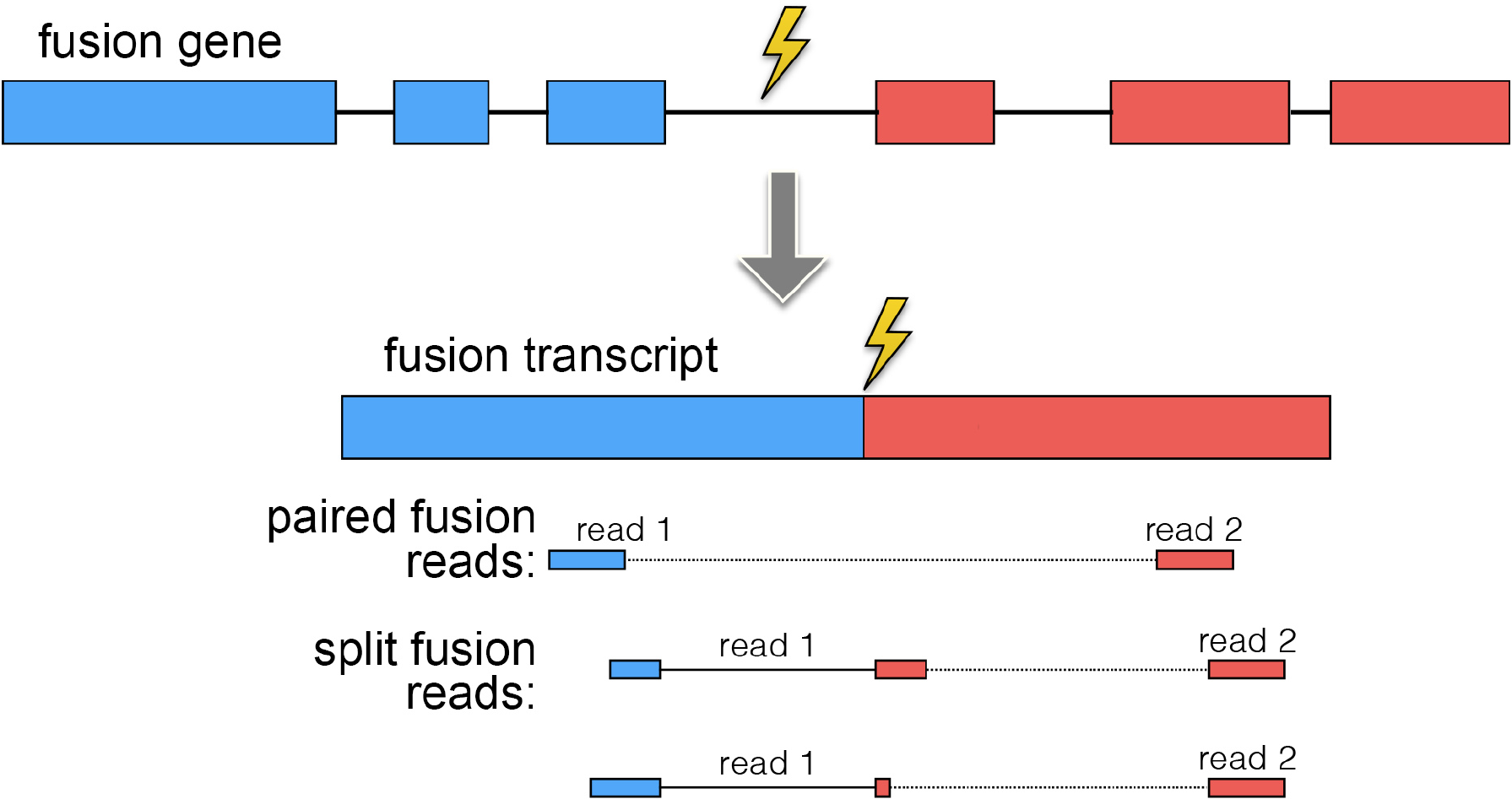
Reads meeting specific criteria are passed to pizzly. A fusion gene resulting from chromosomal rearrangement, is transcribed into a fusion transcript. Standard kallisto does not pseudoalign reads spanning the fusion junction to the transcriptome. In the fusion-modified kallisto, reads meeting specific criteria are reported as candidate fusions for use with pizzly.

### pizzly Fusion Stage

Unlike kallisto, which is genome annotation agnostic, pizzly performs a more detailed analysis of the reads to determine fusion breakpoints by taking the annotation of the transcriptome into account. pizzly accepts the annotation in GTF format, which records the genomic locations of exons as well as functional annotation of genes (e.g. whether they are protein coding).

The input to pizzly is the set of read pairs that kallisto flagged as potentially spanning fusion junctions. In the first step of pizzly, each read is evaluated independently and reads that are classified as false positives are rejected. pizzly uses several criteria to reduce the number of false positives. First, reads that map to transcripts in multiple genomic locations are discarded. Next, since kallisto is unaware of which transcripts belong to which genes, its output can contain false positives resulting from two distinct transcripts associated with the same gene. These are typically from isoforms that were not present in the reference transcriptome and are discarded. The output of kallisto also contains false positives where read pairs originate from a single transcript with mismatches (SNP, etc.) to the reference, but kallisto assigned it to two distinct but similar genes. Rather than relying on annotation of gene families to filter these cases, they are filtered using approximate sequence alignments. Matching *k*-mers from one end are considered and approximate matches to the transcripts of the other end are then examined (Figure 3). Instead of aligning each *k*-mer to the entire transcriptome with allowance for mismatches, only potential candidates kallisto identified for the other end are considered. If any such approximate match is found for either one of the ends, the read pair is discarded.

**Figure 3:**
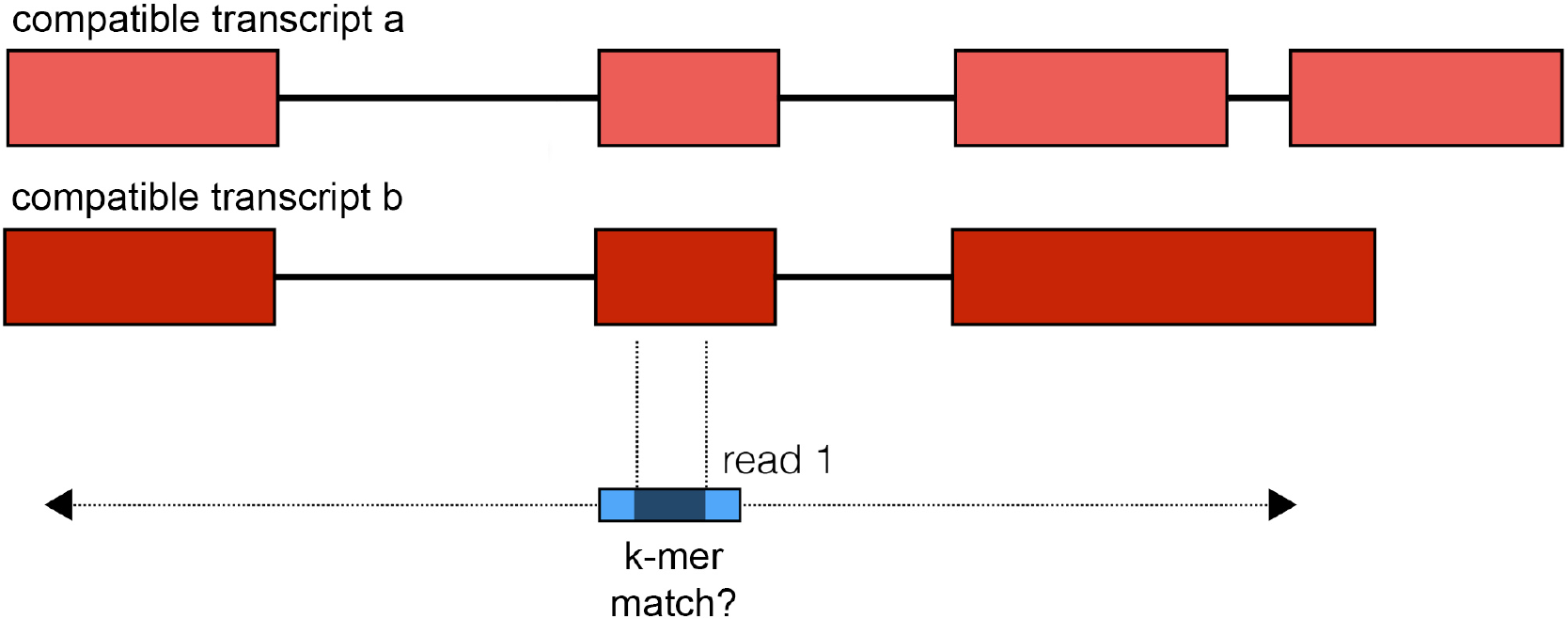
A check for alignment to partner transcript ensures correct fusion. The *k*-mers from the first part of the candidate fusion, the blue “read 1”, are aligned with mismatch allowance to all compatible transcripts from the second part of the candidate fusion. If a match is found, this false positive is discarded. The same is repeated for the other end of the candidate fusion.

Third, the entire read is required to align to the transcripts identified by kallisto. If each read aligns to representative transcript sequences, the read pair is labeled “paired” (Figure 2). In cases where either read cannot be aligned to the potential transcript, an attempt is made to compute a split-read alignment (Figure 4). In order to limit the potential for false positive splits, the search space is restricted to only include split sequences that fit within a specified insert length (Figure 5). Furthermore, the split breakpoint cannot be to close to the end of the read. Reads that can be split-aligned are labeled as “split.”

**Figure 4:**
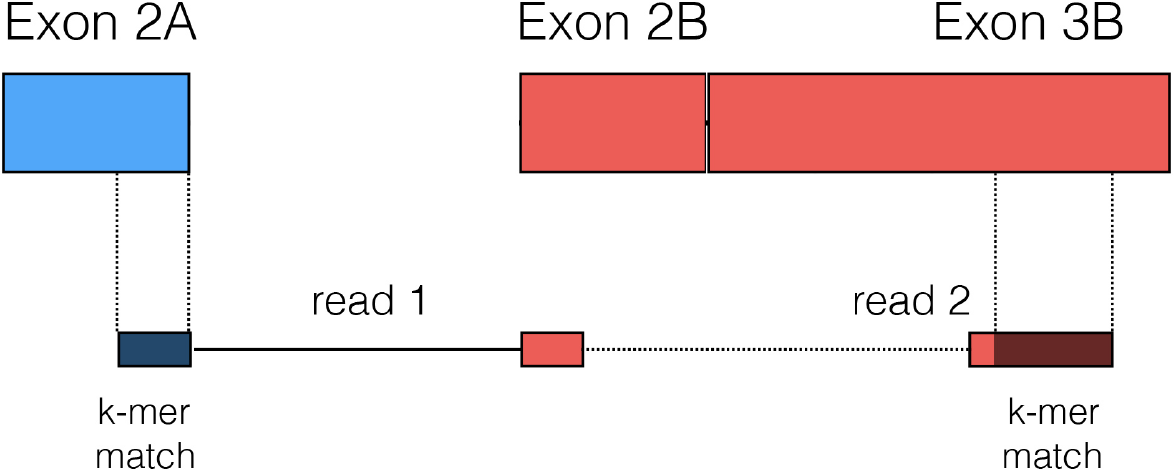
A read can be split-aligned between the two fusion transcripts. Split-read alignments provide breakpoint sequence resolution.

**Figure 5:**
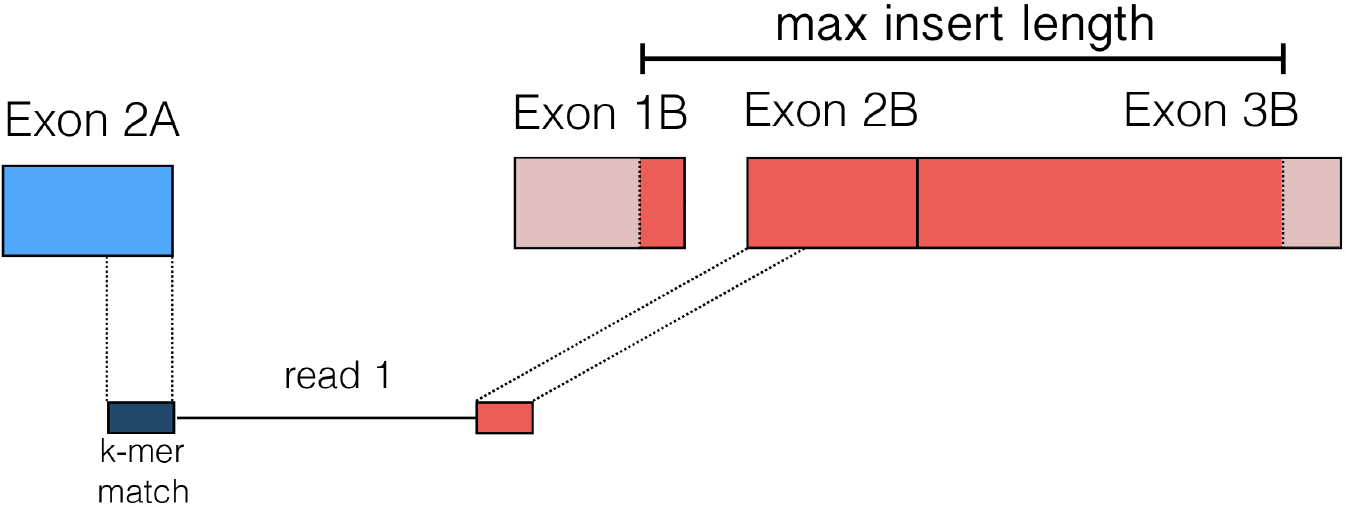
Split alignment search. The search space for split alignments is limited to the matching part of the compatible transcript up to the fragment insert length.

Finally, when all the reads have been filtered, the resultant information is aggregated on a gene-to-gene fusion level. The potential transcript junctions are filtered based on the number of reads supporting the junction. Additionally for split reads, the distance of the breakpoint to exon boundaries is required to be less than 10 bp on both sides. For junctions only supported by pairs, the distance to the nearest internal exon boundaries is required to be consistent with the specified insert length. After filtering pizzly reports the number of paired and split reads supporting the fusion junction. Additionally, pizzly reports each potential transcript junction, the number of read pairs supporting the transcript-level fusion as well as sequence of the fused transcripts and the individual reads supporting the junction.

## Results

We applied our method to multiple datasets to assess the precision/recall and running time of pizzly for fusion detection. To benchmark our performance, we compared our results to existing fusion software packages, using the same datasets to evaluate the performance of each program.

### Datasets

We assessed performance on four types of data that together represent a range of challenges in gene fusion detection.

#### Positive Control

Two positive synthetic datasets of different read lengths were used, both consisting of reads from the same 50 fusion transcript sequences:

a. A positive dataset containing 57,209 75nt paired-end synthetic reads corresponding to 50 fusion transcripts, with 4,300 reads directly covering the fusion junctions. This dataset was created by the FusionMap developers^16^ and used in a previous fusion tool evaluation^11^.
b. A positive dataset containing 200,000 100nt paired-end synthetic reads corresponding to the same 50 fusion transcripts, simulated using the RSEM read simulator^17^.

#### Negative Control

A negative dataset containing 30 million 100nt paired-end synthetic reads simulated using RSEM read simulator using as background expression sample SRR066679 originating from H1 human embryonic stem cells^18^.

#### Real Data with Validation

Fusion detection may perform well on simulated datasets and yet behave differently when presented with real data generated via RNA-Seq. We used real data to assess performance under standard experimental conditions.

An mRNA-Seq dataset, SRR1659964, consisting of 93,867,189 100nt paired-end reads and containing sequence from nine synthetic fusion transcript RNA constructs. This dataset was generated by Tembe et al.^19^ by pooling the nine fusion constructs at eqimolar concentration and adding -6.17 log10pMol to a 1ug aliquot of total RNA, preparing an Illumina TruSeq stranded mRNA library, and sequencing on an Illumina HiSeq 2500.

### Tools for Benchmarking

There are several fusion detection software packages available. We chose to focus our comparison on software that met the following criteria: (1) used in the field (based upon citation number or newness), (2) high performance as evaluated in earlier benchmarking papers^11,20^, and (3) ability to run the software, which is not always a trivial task. The tools meeting these criteria are described in Table 1 below.

**Table 1.** Fusion detection tools compared to pizzly for benchmarking

| tool | version | parameters | reference used |
| --- | --- | --- | --- |
| MapSplice | 2.2.1 | default | ensemble hg38 |
| EricScript | 0.0.5 | default | E.S. formatted hg38 |
| FusionCatcher | 0.99.6a | --skip-banned-fusions, -<br>-skip-known-fusions | F.C. formatted hg38 |
| JAFFA | 1.07 | default: hybrid on 75bp,<br>direct on 100 | UCSC hg38, provided by<br>JAFFA |
| PRADA | 1.2 | suggested -junL 80% of<br>read length | PRADA formatted hg37 |
| STAR-Fusion | 0.8.0 | default | S.F. formatted GRCh38v23 |

### Results

Tools were run using four cores where possible. MapSplice was run on an Amazon Elastic Compute Cloud instance Ubuntu machine type c4.4xl (16cpu, 30GiB memory). PRADA was run with the suggested length of constructed junctions (-junL) of 80% of read length, which for the 75nt dataset was 60 and for the 100nt dataset was 80. FusionCatcher has hardcoded known true and false positive fusions into the program. While hardcoding known fusions may be helpful in a clinical setting, in exploratory analysis, such as is performed in research, this is a limiting restriction that may bias results and reduces the utility of the program. We have therefore bypassed the hardcoded setting in this comparison. JAFFA has three run modes: Assembly, Hybrid, and Direct, which are suggested for different read lengths. For short reads JAFFA assembles the reads to search for fusions in assembled contigs. For longer reads, JAFFA suggests no assembly. We followed the suggested modes for read lengths of 75nt and 100nt.

The results of the tests are summarized in Tables 2-5. PRADA failed to run on all 100nt datasets, finding fusions only for the 75bp dataset. EricScript predicted true fusions decently, but had difficulty filtering out false positives. This would limit its utility in exploratory analyses. JAFFA performed well with clean, simulated datasets but dropped in performance when presented with real datasets, picking up many false positives and missing true fusions. STAR-Fusion performed fairly well, with a fair sensitivity and fair specificity across data types. All of the comparison programs required a long time to run, ranging on the larger spike-in dataset from over 26 hours for MapSplice to 1 hour 18 minutes for the fastest, STAR-Fusion.

**Table 2.** **75nt positive control** pizzly detects the most fusions. The starred “false positives” reported by pizzly were filtered out in future pizzly iterations using our updated method discussed below. Timing is based on using 4 cores where possible.

| tool | true positive | false positive | sensitivity % | clock time |
| --- | --- | --- | --- | --- |
| MapSplice | 42 | 1 | 84 | 2m1s |
| EricScript | 39 | 0 | 78 | 9m34s |
| FusionCatcher | 31 | 0 | 62 | 5m33s |
| JAFFA | 43 | 0 | 86 | 3m56s |
| PRADA | 32 | 0 | 64 | 4m5s |
| STAR-Fusion | 45 | 1 | 90 | 6m59s |
| pizzly | 45 | 2* | 90 | 1m8s |

**Table 3.** **100nt positive control**

| tool | true positive | false positive | sensitivity % |
| --- | --- | --- | --- |
| MapSplice | 39 | 1 | 78 |
| EricScript | 41 | 1 | 82 |
| FusionCatcher | 41 | 0 | 82 |
| JAFFA | 47 | 0 | 94 |
| PRADA | 0 | 0 | 0 |
| STAR-Fusion | 41 | 0 | 82 |
| pizzly | 46 | 2* | 92 |

**Table 4.** **100nt negative control** All tools but EricScript perform well, picking up only minor numbers of false positives.

| tool | false positive |
| --- | --- |
| MapSplice | 3 |
| EricScript | 82 |
| FusionCatcher | 2 |
| JAFFA | 1 |
| PRADA | 1 |
| STAR-Fusion | 0 |
| pizzly | 3 |

**Table 5.** **Spike-in real dataset containing nine true synthetic fusions** Since this is a real dataset, “false positives” has been replaced with “other” to allow for possible real fusions. In the study, only one other fusion was confirmed in the background RNA sample. Timing is based on using 4 cores where possible.

| tool | true (9) | other | clock time |
| --- | --- | --- | --- |
| MapSplice | 7 | 2 | 26h 37m |
| EricScript | 9 | 575 | 10h 17m |
| FusionCatcher | 9 | 11 | 5h 37m |
| JAFFA | 7 | 458 | 20 17m |
| PRADA | 0 | 0 | 19h 7m |
| STAR-Fusion | 8 | 5 | 1h 18m |
| pizzly | 9 | 624 | 8.5m |

pizzly showed high sensitivity across all datasets and high specificity in the simulated datasets. pizzly runs much faster than any of the other programs, taking just 8.5 minutes on the large spike-in dataset. Since kallisto-pizzly runs in such a short amount of time, pizzly offers the possibility for iteratively refining filtering strategies based on analyses of pizzly output and biologically relevant heuristics. This could be applied in the future to further improve pizzly’s specificity.

## Discussion

Our results show that pizzly is highly accurate in a range of benchmarks, positive and negative controls. The program is also fast, making it possible to reproducibly and consistently annotate fusions in large cancer datasets. Furthermore, the speed of pizzly enables exploration of different filters, augmented transcriptomes to be tested and can assure robustness of results with tests using a range of filter parameters. There is also room for improvement of pizzly as the biology of fusions is better understood.

For example, in the 75nt positive control, pizzly reported two false positives (starred in Table 2). Upon further investigation, we found that these false fusions originated from genes which overlap in the genome. In the first case pizzly predicts the “true” fusion PLEKHO2-KIF4A and the “false” fusion AC069368.3-KIF4A, however AC069368.3 is made up of the exons of PLEKHO2 and the neighboring ANKDD1A gene and except for an alternative start site the sequences of these predicted fusions are identical. In the second case pizzly predicts the “true” fusion BSG-COX6A1 and the “false” fusion BSG-AL021546.6 and identifies sixteen possible transcript sequences arising from the eight BSG transcripts and the two other transcripts. The original transcript estimates from kallisto predicted that only two of the BSG transcripts are present in the sample and only COX6A1 and not AL021546.6 (Table 6). Given the breakpoint sequences of these sixteen possible fusion transcripts, we built a new kallisto index with the added potential fusions and re-quantified the original reads with kallisto using the updated index (Table 7).

**Table 6.** **Original kallisto read counts for suspected fusion transcripts**

| transcript | estimated reads |
| --- | --- |
| COX6A1 | 3002 |
| AL021546.6 | 0 |
| BSG-202 | 2218 |
| BSG-204 | 4 |

**Table 7.**
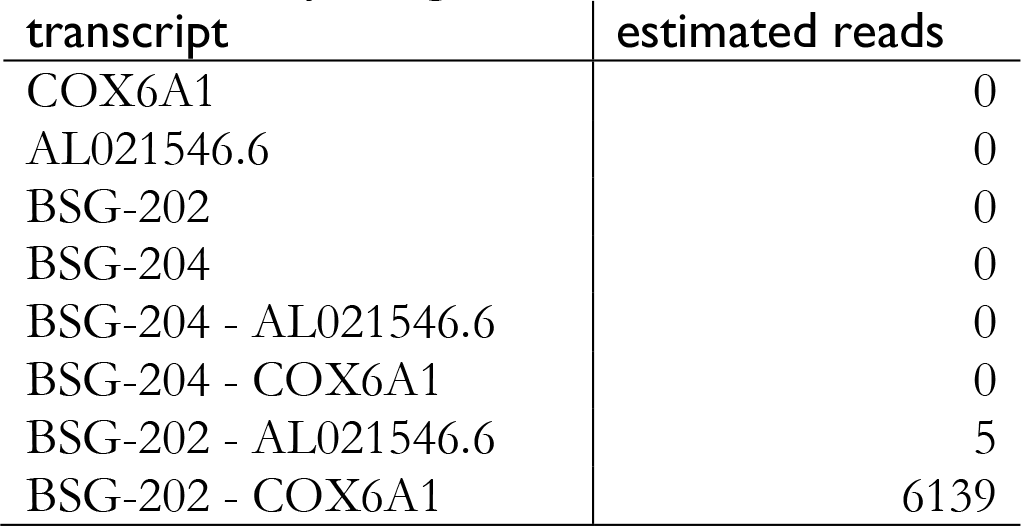
**kallisto is able to correctly assign reads to fusions at the transcript level**

| transcript | estimated reads |
| --- | --- |
| COX6A1 | 0 |
| AL021546.6 | 0 |
| BSG-202 | 0 |
| BSG-204 | 0 |
| BSG-204 - AL021546.6 | 0 |
| BSG-204 - COX6A1 | 0 |
| BSG-202 - AL021546.6 | 5 |
| BSG-202 - COX6A1 | 6139 |

By adding a second kallisto step after fusion transcript assembly, we were able to increase detection of fusion transcripts while reducing false positives. In addition, this extra kallisto step allowed us to estimate transcript fusion abundance. While all other fusion detection programs report transcript abundance only on the level of read counts, we were able to output the more meaningful and accurate transcript per million (TPM) report for detected fusions. To test the accuracy of TPM estimation for fusion transcripts, we applied the kallisto-pizzly-kallisto method to the spike-in dataset that contains fusion transcripts aliquoted at a known concentration. True fusion transcript abundance was calculated as follows: starting with 1ug of RNA, with mRNA being 1-5% of the total and each transcript being on average 2,500nt, the concentration of background RNA is approximately 1.17pmol and mRNA approximately 0.058pmol. The spike-in was aliquoted at -6.17 log10pmol. Under the assumption that the mRNA is between 1-5% of the total RNA, the total spike-in concentration should be 11-60 TPM divided across the nine synthetic fusions. kallisto’s estimated fusion abundances add up to 31.6TPM, which is within the expected range (Table 8). Thus kallisto should be valuable not only for identifying fusions, but for associating their expression with cancer phenotypes and outcomes.

**Table 8.** **kallisto reported TPM for the nine synthetic fusion transcripts**

| fusion | TPM |
| --- | --- |
| EWSR1-ATF1 | 2.09 |
| TMPRSS2-ETV1 | 8.10 |
| EWSR1-FLI1 | 1.45 |
| NTRK3-ETV6 | 5.12 |
| CD74-ROS1 | 5.37 |
| HOOK3-RET | 3.74 |
| EML4-ALK | 1.31 |
| AKAP9-BRAF | 3.04 |
| BRD4-NUT | 1.35 |

## Acknowledgements

The authors would like to thank David P. Kreil and Paweł P. Łabaj for helpful feedback.

